# Dendritic cell deficiencies persist seven months after SARS-CoV-2 infection

**DOI:** 10.1101/2021.03.18.436001

**Authors:** A Perez-Gomez, J Vitalle, MC Gasca-Capote, A Gutierrez-Valencia, M Trujillo-Rodriguez, A Serna-Gallego, E Muñoz-Muela, MR Jimenez-Leon, M Rafii-El-Idrissi Benhnia, I Rivas-Jeremias, C Sotomayor, C Roca-Oporto, N Espinosa, C Infante-Dominguez, JC Crespo-Rivas, A Fernández-Villar, A Pérez-González, LF Lopez-Cortes, E Poveda, E Ruiz-Mateos, on behalf of COHVID-GS and Virgen del Rocío Hospital COVID-19 working team

## Abstract

Severe Acute Respiratory Syndrome Coronavirus (SARS-CoV)-2 infection induces an exacerbated inflammation driven by innate immunity components. Dendritic cells (DCs) play a key role in the defense against viral infections, for instance plasmacytoid DCs (pDCs), have the capacity to produce vast amounts of interferon-alpha (IFN-α). In COVID-19 there is a deficit in DC numbers and IFN-α production, which has been associated with disease severity. In this work, we described that in addition to the DC deficiency, several DC activation and homing markers were altered in acute COVID-19 patients, which were associated with multiple inflammatory markers. Remarkably, previously hospitalized and non-hospitalized patients remained with decreased numbers of CD1c+ myeloid DCs and pDCs seven months after SARS-CoV-2 infection. Moreover, the expression of DC markers as CD86 and CD4 were only restored in previously non-hospitalized patients while integrin β7 and indoleamine 2,3- dyoxigenase (IDO) no restoration was observed. These findings contribute to a better understanding of the immunological sequelae of COVID-19.

## INTRODUCTION

Coronavirus disease 19 (COVID-19) is caused by severe acute respiratory syndrome coronavirus 2 (SARS-CoV-2) infection and may progress with mild symptoms or asymptomatically in most of the individuals, while others experience an acute respiratory distress syndrome (ARDS) and poorer prognosis, including death (Guan et al., 2020). Disease severity depends on the balance between host immune response, viral replication and tissue and organ damage. In severe COVID-19 there is a deregulation of this response, characterized by an hyper-inflammation driven by innate immunity, characterized by very high levels of cytokines and pro-inflammatory biomarkers, also known as cytokine storm (Chi et al., 2020; Zhou et al., 2020b).

One of the innate immune cell types that may play a pivotal role in the response against SARS-CoV-2 are the dendritic cells (DCs). There are two main DC types, conventional or myeloid DCs (mDCs) which include CD1c+, CD16+ and CD141+ mDC subsets, and plasmacytoid dendritic cells (pDCs). In general, DCs participate in antigen presentation, cytokine production, control of inflammatory responses, tolerance induction, immune cell recruitment, and viral dissemination. However, the role of these cells in response to acute SARS-CoV-2 infection and the recovery in convalescent subjects is not fully characterized. Some studies have shown a decrease of DC numbers in response to infection in peripheral blood (Kvedaraite et al., 2021) and also an association with disease severity (Zhou et al., 2020a). This deficiency seems to be due to the migration of some DC subsets, such as CD1c, to the lung (Sánchez-Cerrillo et al., 2020), and probably to other inflammatory foci. pDCs also seems to play a key role in COVID-19 (Arunachalam et al., 2020). pDCs are the main type I interferon (IFN-I) producers, with 1000-fold production compared to other immune cell types (Swiecki and Colonna, 2010). IFN-I is known to have an essential role in viral infections (Muller et al., 1994). Significantly, pDCs depletion has been associated with poor COVID-19 prognosis (Laing et al., 2020). Moreover, critical patients showed a highly impaired IFN-I response (Arunachalam et al., 2020) associated with high viral load and aggravated inflammatory response (Hadjadj et al., 2020).

The recovery of DC defects after COVID-19 could be crucial, since the normalization of the innate immune system after the acute insult would mean the system’s readiness to respond to new viral and bacterial challenges. However, the recovery of DC cell numbers and function after COVID-19 is unknown. This recovery is also important in the sense that a variable proportion of people who have overcome COVID-19 show clinical sequelae (Carfì et al., 2020) which relation with innate immune defects needs to be clarified. Thus, the aim of the study was to analyze DC defects associated with SARS-CoV-2 infection, COVID-19 severity and whether these defects were restored after a median of seven months after the resolution of the infection.

## RESULTS

### Patients with acute SARS-CoV-2 infection show a considerable decrease in DC percentages and TLR9-dependent IFN-α production

In order to investigate the effect of SARS-CoV-2 infection on the innate immune system, we first analyzed the percentages of total DCs and the different subsets in acute SARS-CoV-2 infected patients (COVID-19 patients) compared with age and sex matched healthy donors (HD). Specifically, we measured mDCs (CD123- CD11c+), including CD1c+, CD16+ and CD141+ mDC subsets, and pDCs (CD123+ CD11c-) (Figure S1 A). Our results showed that acute COVID- 19 patients exhibited a significant decrease in the percentages of total mDCs mainly due to CD1c+ mDCs decreased in comparison with HD. Meanwhile CD16+ and CD141+ mDCs remained at similar levels of HD (Figure 1A). Remarkably, the percentage of pDCs in acute COVID-19 patients was considerably diminished with respect to HD (Figure 1B left). Then, we calculated the ratio mDC/pDC in the different subjects, which was much lower in HD that in COVID-19 patients (Figure S2). pDCs are known to be the main producers of IFN-α (Swiecki and Colonna, 2010). Therefore, to study their function in SARS-CoV-2 infection, we stimulated peripheral blood mononuclear cells (PBMCs) with CpG oligodeoxynucleotides class A (CpG)-A, a Toll-like receptor (TLR)-9 dependent stimulation, and we analyzed IFN-α production. We found that IFN-α production in acute COVID-19 was much lower than in HD (Figure 1B right). To clarify if the decreased IFN-α production was due to a diminished percentage of pDCs, we performed a correlation analysis and we found that the IFN-α production was positively associated with the percentage of pDCs in both acute COVID-19 patients and HD (Figure 1C). Although at lower levels, mDCs can also produce IFN (Jongbloed et al., 2010), so we also correlated both parameters and we found that only CD141+ mDCs showed a slight correlation with the secretion of this cytokine (Figure S3). In conclusion, patients with acute SARS-CoV-2 infection exhibit a deficit in DC numbers and also decreased TLR9-dependent IFN-α production.

**Figure 1.**
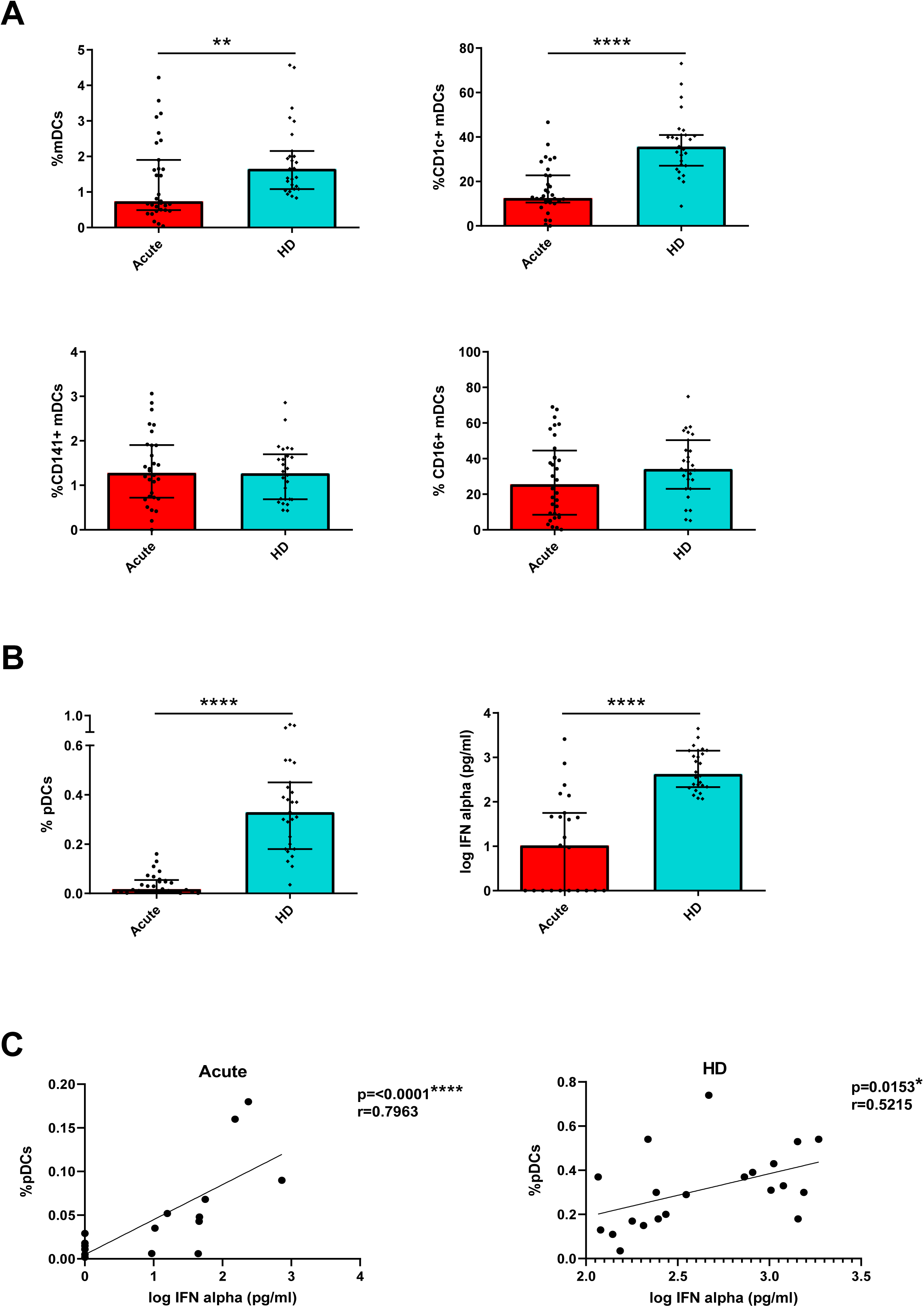
Patients with acute SARS-CoV-2 infection show a considerable decrease in DC percentages and TLR9-dependent IFN-α production. Bar graphs representing the percentage of total mDCs, CD1c+, CD141+ and CD16+ mDCs (A) and the percentage of pDCs and IFN-α production in response to CpG-A (B) in acute SARS-CoV-2 infected patients (acute) and healthy donors (HD). The median with the interquartile range is shown. Please see Figure S1 for the gating strategy and Figure S2 for the mDC/pDC ratio. (C) Correlation between the percentage of pDCs and IFN-α production in acute patients and HD. See Figure S3 for correlations of mDC percentages. Each dot represents an individual. *p < 0.05, **p < 0.01, ***p < 0.001, ****p < 0.0001. Mann-Whitney U test was used for groups’ comparisons and Spearman test for non-parametric correlations.

### Acute SARS-CoV-2 infected patients show an altered pattern of DC activation markers

Afterwards, we analyzed the expression of DC activation markers in acute COVID-19 patients and HD. We measured the expression of homing receptors ((integrin-β7 (β7) and C-C chemokine receptor type 7 (CCR7)), co-stimulatory molecules (CD86 and CD4), and markers of immune tolerance and suppression ((Indoleamine 2,3-dioxygenase (IDO) and Programmed Death-ligand 1 (PD-L1)) (Figure S1 B). Most of the DC subpopulations, presented lower percentage of β7, specially total mDCs, CD1c+ mDCs and pDCs and a higher percentage of CCR7+ DCs in acute COVID-19 patients compared with HD (Table 1). We also found lower percentage of CD86+ cells in acute patients in CD1c+ and CD16+ mDCs and pDCs. No differences were in CD4+ DC levels (Table 1). Lastly, acute COVID-19 patients showed higher percentage of IDO+ cells within CD1c+ and CD16+ mDCs compared with HD, while a lower percentage PD-L1+ was seen within pDCs (Table 1). These results are indicative of alterations in different homing and activation patterns of DCs in response to SARS-CoV-2 infection.

**Table 1.**
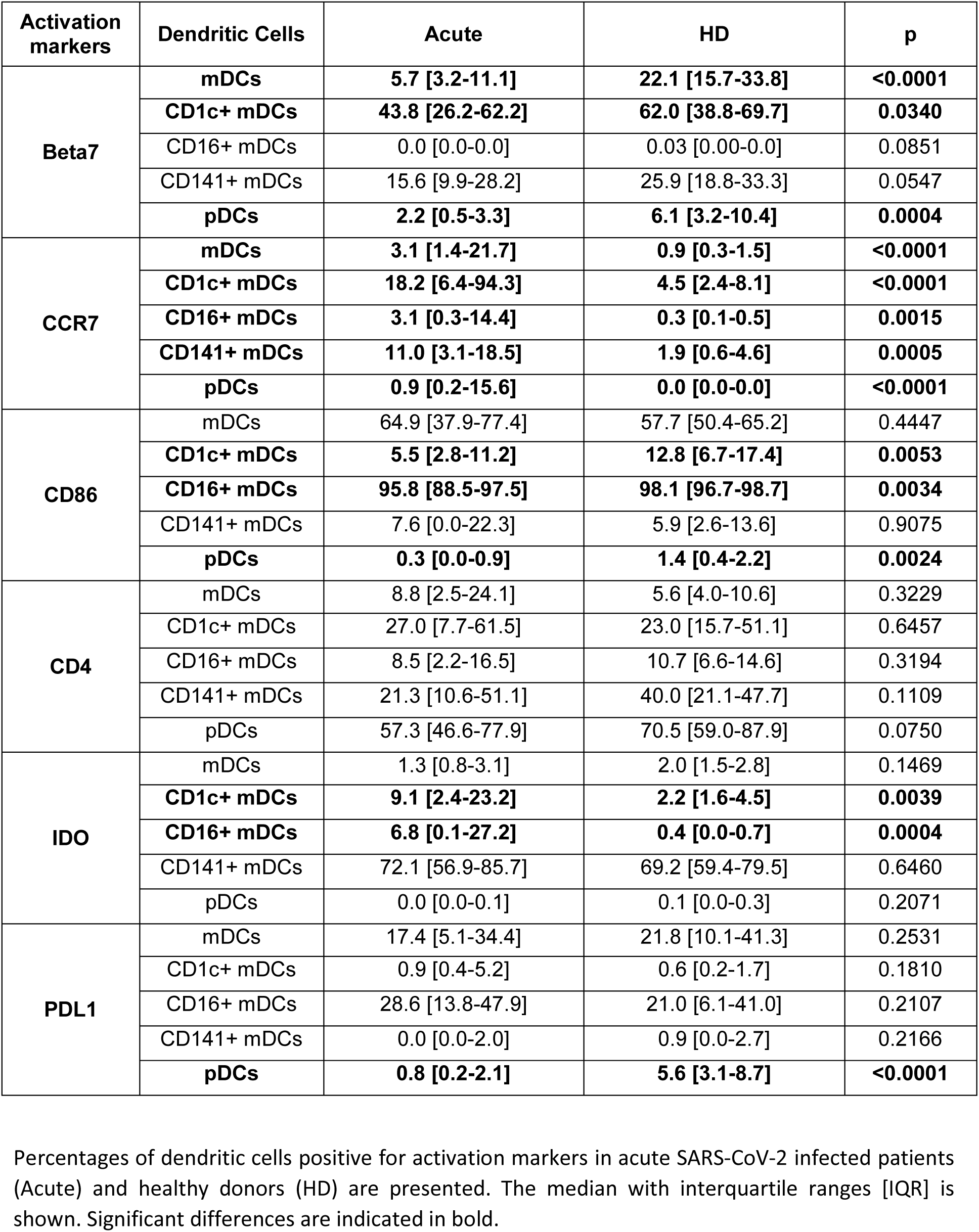
Acute SARS-CoV-2 infected patients show an altered pattern of DC homing and activation markers

| Activation markers | Dendritic Cells | Acute | HD | p |
| --- | --- | --- | --- | --- |
| **Beta7** | mDCs | **5.7 [3.2-11.1]** | **22.1 [15.7-33.8]** | **<0.0001** |
|  | **CD1c+ mDCs** | **43.8 [26.2-62.2]** | **62.0 [38.8-69.7]** | **0.0340** |
|  | CD16+ mDCs | 0.0 [0.0-0.0] | 0.03 [0.00-0.0] | 0.0851 |
|  | CD141+ mDCs | 15.6 [9.9-28.2] | 25.9 [18.8-33.3] | 0.0547 |
|  | **pDCs** | **2.2 [0.5-3.3]** | **6.1 [3.2-10.4]** | **0.0004** |
| **CCR7** | mDCs | **3.1 [1.4-21.7]** | **0.9 [0.3-1.5]** | **<0.0001** |
|  | **CD1c+ mDCs** | **18.2 [6.4-94.3]** | **4.5 [2.4-8.1]** | **<0.0001** |
|  | **CD16+ mDCs** | **3.1 [0.3-14.4]** | **0.3 [0.1-0.5]** | **0.0015** |
|  | **CD141+ mDCs** | **11.0 [3.1-18.5]** | **1.9 [0.6-4.6]** | **0.0005** |
|  | **pDCs** | **0.9 [0.2-15.6]** | **0.0 [0.0-0.0]** | **<0.0001** |
| **CD86** | mDCs | 64.9 [37.9-77.4] | 57.7 [50.4-65.2] | 0.4447 |
|  | **CD1c+ mDCs** | **5.5 [2.8-11.2]** | **12.8 [6.7-17.4]** | **0.0053** |
|  | **CD16+ mDCs** | **95.8 [88.5-97.5]** | **98.1 [96.7-98.7]** | **0.0034** |
|  | CD141+ mDCs | 7.6 [0.0-22.3] | 5.9 [2.6-13.6] | 0.9075 |
|  | **pDCs** | **0.3 [0.0-0.9]** | **1.4 [0.4-2.2]** | **0.0024** |
| **CD4** | mDCs | 8.8 [2.5-24.1] | 5.6 [4.0-10.6] | 0.3229 |
|  | CD1c+ mDCs | 27.0 [7.7-61.5] | 23.0 [15.7-51.1] | 0.6457 |
|  | CD16+ mDCs | 8.5 [2.2-16.5] | 10.7 [6.6-14.6] | 0.3194 |
|  | CD141+ mDCs | 21.3 [10.6-51.1] | 40.0 [21.1-47.7] | 0.1109 |
|  | pDCs | 57.3 [46.6-77.9] | 70.5 [59.0-87.9] | 0.0750 |
| **IDO** | mDCs | 1.3 [0.8-3.1] | 2.0 [1.5-2.8] | 0.1469 |
|  | **CD1c+ mDCs** | **9.1 [2.4-23.2]** | **2.2 [1.6-4.5]** | **0.0039** |
|  | **CD16+ mDCs** | **6.8 [0.1-27.2]** | **0.4 [0.0-0.7]** | **0.0004** |
|  | CD141+ mDCs | 72.1 [56.9-85.7] | 69.2 [59.4-79.5] | 0.6460 |
|  | pDCs | 0.0 [0.0-0.1] | 0.1 [0.0-0.3] | 0.2071 |
| **PDL1** | mDCs | 17.4 [5.1-34.4] | 21.8 [10.1-41.3] | 0.2531 |
|  | CD1c+ mDCs | 0.9 [0.4-5.2] | 0.6 [0.2-1.7] | 0.1810 |
|  | CD16+ mDCs | 28.6 [13.8-47.9] | 21.0 [6.1-41.0] | 0.2107 |
|  | CD141+ mDCs | 0.0 [0.0-2.0] | 0.9 [0.0-2.7] | 0.2166 |
|  | **pDCs** | **0.8 [0.2-2.1]** | **5.6 [3.1-8.7]** | **<0.0001** |

Percentages of dendritic cells positive for activation markers in acute SARS-CoV-2 infected patients (Acute) and healthy donors (HD) are presented. The median with interquartile ranges [IQR] is shown. Significant differences are indicated in bold.

### IFN-α production is associated with COVID-19 severity

The next step of this study was to investigate whether DC numbers and their function might be different in acute COVID-19 depending on disease severity. Therefore, we classified acute COVID-19 patients in two groups: severe ((high oxygen support requirement and Intensive Care Unit (ICU) admission or death)) and mild (low oxygen requirement and no ICU admission) (Table S1). Our results did not show any significant difference in the percentage of mDCs and subpopulations between severe and mild COVID-19 patients (Figure 2A). However, we found increased levels of total CCR7+ mDCs and PD-L1+ CD141+ mDCs in severe patients (Figure S4). Regarding pDCs, again we did not observe significant differences, although there was a trend to lower levels in severe than mild patients (Figure 2B, left). Importantly, we did find a considerable decrease in TLR9-dependent IFN-α production in severe subjects compared to mild patients (Figure 2B, right). In summary, acute SARS-CoV-2 infected patients with severe symptoms exhibit a lower capacity to produce IFN- α than patients with mild symptoms.

**Figure 2.**
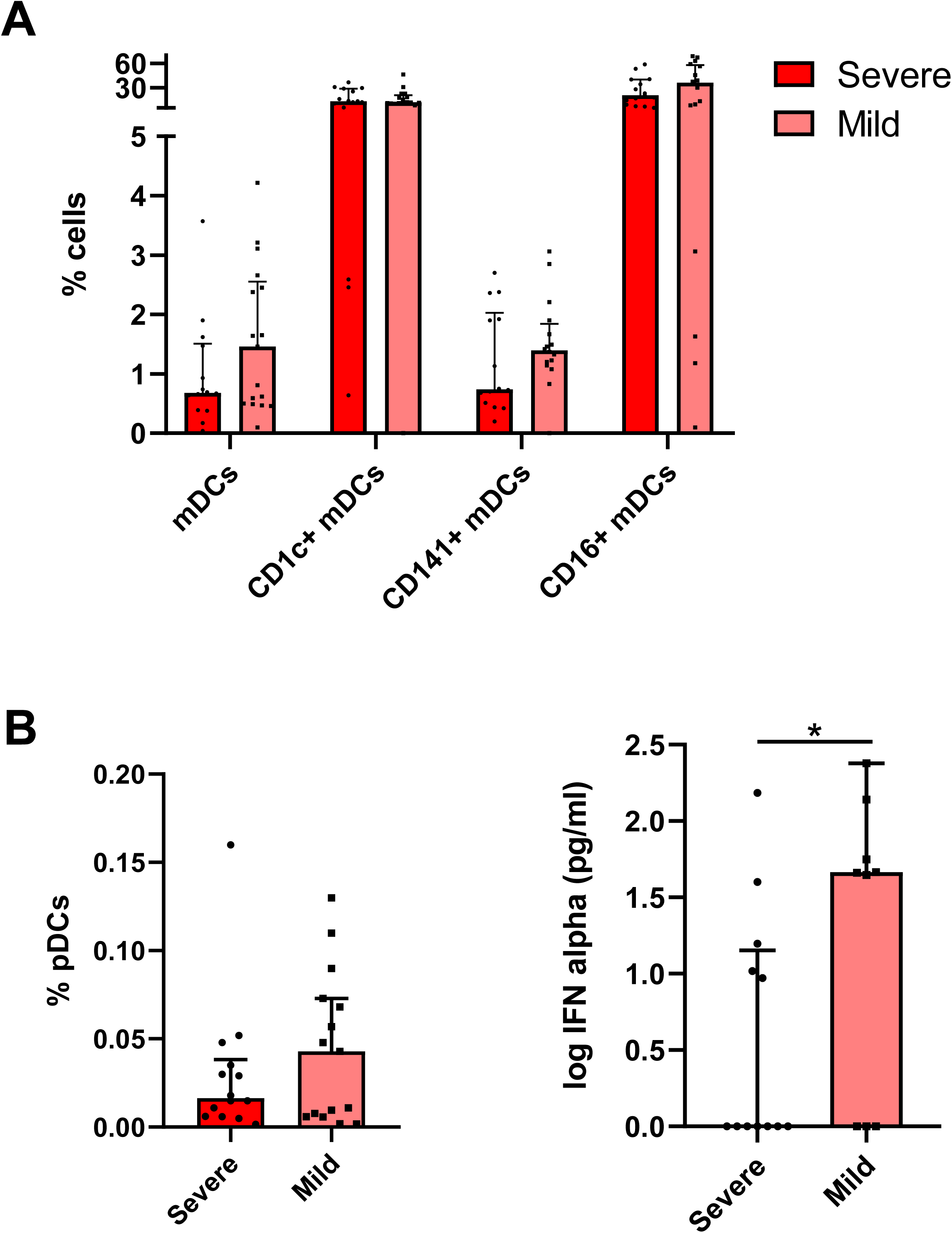
IFN-α production is associated with COVID-19 severity. Bar graphs representing the percentage of total mDCs, CD1c+, CD141+ and CD16+ mDCs subsets (A) and the percentage of pDCs and IFN-α production in response to CpG-A (B) in acute severe and mild SARS-CoV-2 infected patients. See Figure S4 for the percentage of CCR7+ and PD-L1+ DCs in severe and mild acute SARS-CoV-2 infected patients. The median with the interquartile range is shown and each dot represents an individual. *p < 0.05. Mann-Whitney U test was used for groups’ comparisons.

### DC parameters are differentially associated to inflammation markers in mild and severe acute SARS-CoV-2 infected patients

Then, DC numbers and activation markers were correlated to multiple inflammatory marker levels, including clinical biomarkers ((C-reactive protein (CRP), D-dimers and lactate dehydrogenase (LDH)), pro-inflammatory cytokines ((tumor necrosis factor (TNF)-α, interleukin (IL)-6, IL-8, IL1-β, macrophage inflammatory protein (MIP1)-α, MIP1-β, interferon inducible protein (IP)-10 and interferon (IFN)-γ) and soluble (sCD25)), and neutrophil numbers. These correlations were done in the overall group of patients during acute infection and also dividing in both severe and mild COVID-19. In the overall population, we observed correlations of dendritic cell subset levels with different pro-inflammatory cytokines and clinical biomarker levels (Figure S5). Interestingly, we observed a different correlation pattern in severe and mild patients and of note, more associations were found in mild patients (Figure 3). On one hand, regarding COVID-19 patients with mild symptoms, the percentages of DC subpopulations were inversely correlated with D-dimers, IL- 6, IL-8, sCD25 levels and neutrophil numbers, while they were positively correlated with TNF-α, IL-1β, MIP-1α, MIP1-β and IFN-γ levels, with the exception of CD16+ mDCs that were negatively correlated with most of the inflammatory parameters. It is remarkable, that the percentage of pDCs showed a strong inverse correlation with D-dimer levels and neutrophil numbers. Focusing on DC homing and activation markers, regarding the expression of β7 in DCs, inverse associations prevailed, highlighting the strong correlations found in CD16+ β7+ mDCs with D-dimers and in β7+ pDCs with IL1-β. In contrast, the expression of CD86 and IDO in DCs was predominantly positively associated to several inflammatory markers, mainly in the case of CD141+ mDCs and pDCs (Figure 3A). On the other hand, in severe COVID-19, many associations were lost (e.g. IDO expression) and others were opposite (e.g. CD86), comparing with mild patients. For instance, remarkably, the DC percentages and the expression of β7 and CD86, the associations found with inflammatory marker levels showed an opposite trend. (Figure 3B). Therefore, we conclude that DC levels and activation markers are associated to the inflammatory status of acute SARS-CoV-2 infected subjects, with a differential profile between patients with severe symptoms compared to those with mild symptoms.

**Figure 3.**
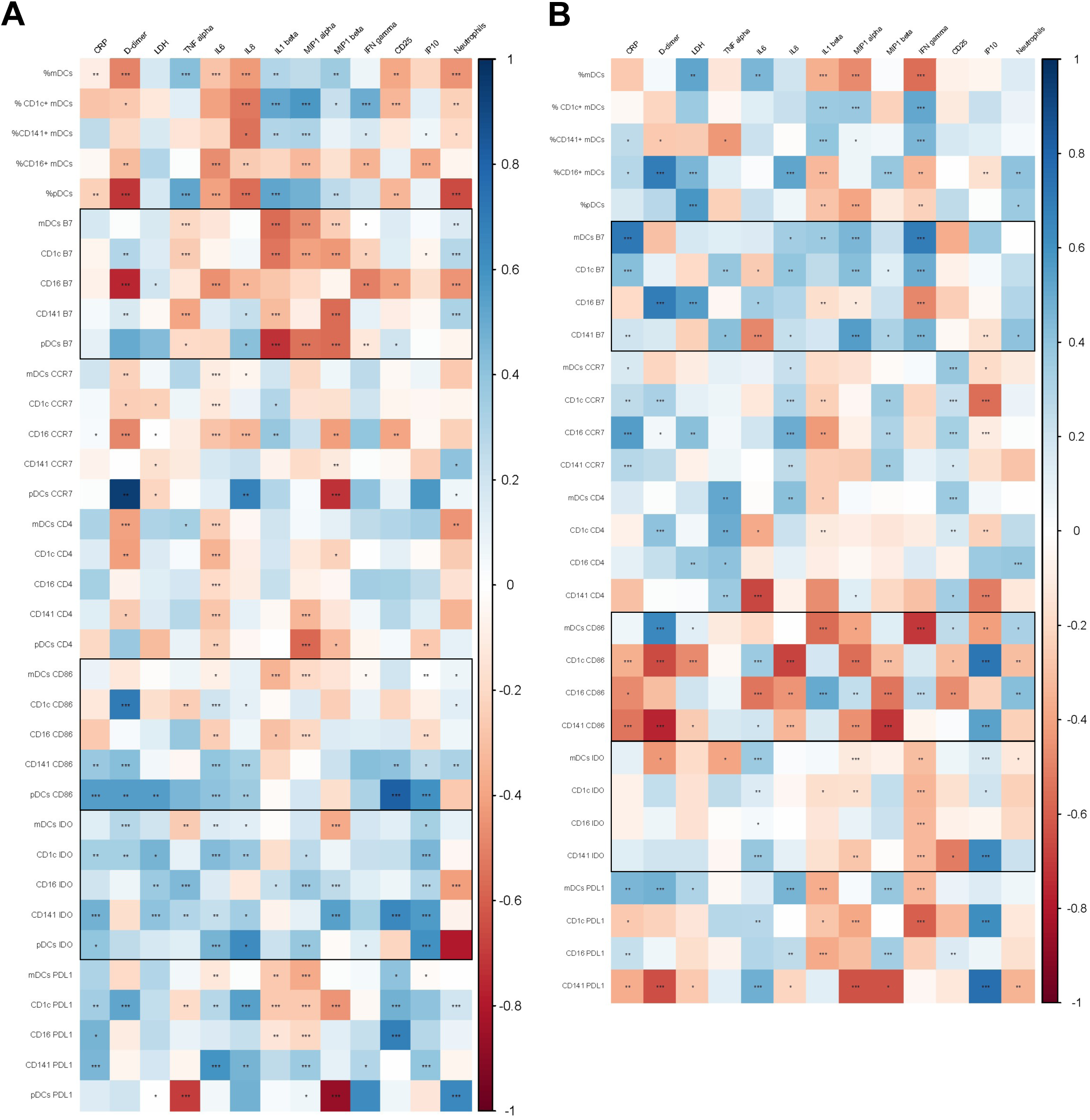
DC parameters are differentially associated to inflammatory markers in mild and severe acute SARS-CoV-2 infected patients. Heatmap graphs representing correlations between the percentages of DC subpopulations and the percentages of DCs expressing activation and homing markers with inflammatory marker levels including CRP, D-dimer, LDH, TNF-α, IL-6, IL-8, IL1-β, MIP1-α, MIP1-β, IFN-γ, CD25, IP-10 and neutrophil numbers, in mild (A) and severe (B) SARS-CoV-2 infected patients. Blue color represents positive correlations and red color shows negative correlations. The intensity of the color indicates the R coefficient. *p < 0.05, **p < 0.01, ***p < 0.001. Please see Figure S5 for correlations in all acute patients. Spearman test was used for non-parametric correlations.

### CD1c+ mDC and pDC levels are not normalized seven months after SARS-CoV-2 infection

Apart from COVID-19 patients in acute phase, we also studied patients after seven months of SARS-CoV-2 infection (median 208 interquartile range [IQR] [189 – 230]) days after symptoms’ onset, Table S1). Some of these patients were hospitalized during acute infection (Hosp 6M), while others were not (No Hosp 6M). We analyzed the percentages of DC subpopulations in these two groups and compared with HD’s levels. First, we observed a higher percentage of total mDCs on previously hospitalized patients compared with HD (Figure 4A). Regarding mDC subpopulations, while the percentages of CD141+ and CD16+ were not altered, the percentage of CD1c+ mDCs remained lower in patients after seven months compared with HD (Figure 4B-D). Remarkably, the percentage of pDCs also persisted very low and was not restored seven months after the infection in these both groups (Figure 4E), confirmed by the mDC/pDC ratio (Figure 4F). Next, to corroborate that our results were reproducible applying a paired analysis, we studied DC kinetic in a subgroup of subjects with available paired samples, analyzing the percentages of DC subpopulations in the acute phase, 6-8 months later and comparing them with HD. Even though the sample size was lower because of the sample availability, these results reproduced the analysis with unpaired samples (Figure S6). When we measured the TLR9-dependent IFN-α production in hospitalized and non-hospitalized patients seven months after the infection, we did not find any significant difference compared to HD (Figure S7). Nevertheless, the percentage of individuals that did not produce IFN-α was higher in hospitalized patients than in non-hospitalized ones or HD (Hosp 6M: 28.5%, No Hosp 6M: 0%, HD: 0%; p=0.05). Here, we conclude that the deficit of CD1c+ mDCs and pDCs is maintained seven months after SARS-CoV-2 infection independently of whether the patients were or not previously hospitalized.

**Figure 4.**
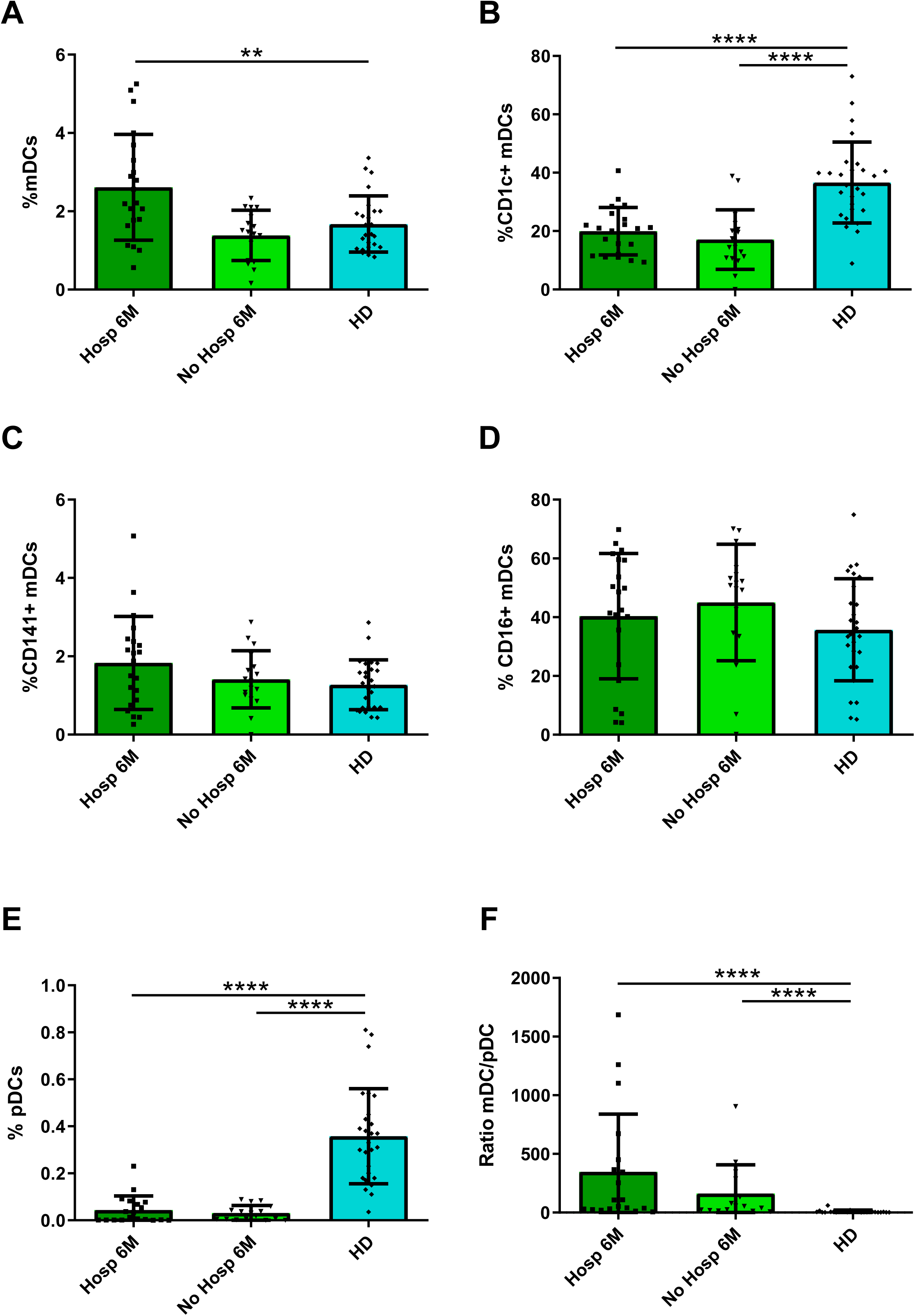
CD1c+ mDC and pDC levels are not normalized seven months after SARS-CoV-2 infection. Bar graphs representing the percentage of total mDCs, CD1c+, CD141+ and CD16+ mDCs, pDCs (A - E) and the DC/pDC ratio (F) in previously hospitalized (Hosp 6M) or previously non-hospitalized (No Hosp 6M) patients seven months after SARS-CoV-2 infection and in healthy donors (HD). The median with the interquartile range is shown and each dot represents an individual. **p < 0.01, ****p < 0.0001. See Figure S6 for the paired analysis of infected patients in acute phase and seven months after infection. Please also see Figure S7 for the IFN-α production in patients seven months after infection. Mann-Whitney U test was used for groups’ comparisons.

### Some DC activation markers are not normalized in previously hospitalized patients seven months after SARS-CoV-2 infection

Afterwards, we measured the DC activation and homing markers in previously hospitalized and non-hospitalized patients seven months after infection, and we compared them with the ones from HD. We observed that the expression of CD86 was lower in CD16+ and CD1c+ mDC subsets from hospitalized patients than in non-hospitalized ones and HD (Figure 5A-B). Similar results were found in the expression of PD-L1 in total mDCs (Figure 5C). Furthermore, hospitalized patients also showed lower levels of CD4 in total mDCs, CD1c+ and CD141+ mDCs and pDCs (Figure 5D-G). In contrast, pDCs from hospitalized patients exhibited higher percentage of CCR7+ cells within pDCs compared with non-hospitalized ones and HD (Figure 5H). In summary, these results show a recovery of some DC activation markers, mainly CD86 and CD4, only in previously non-hospitalized patients, while in more severe patients who required hospitalization, the defects in these markers persisted seven months after infection.

**Figure 5.**
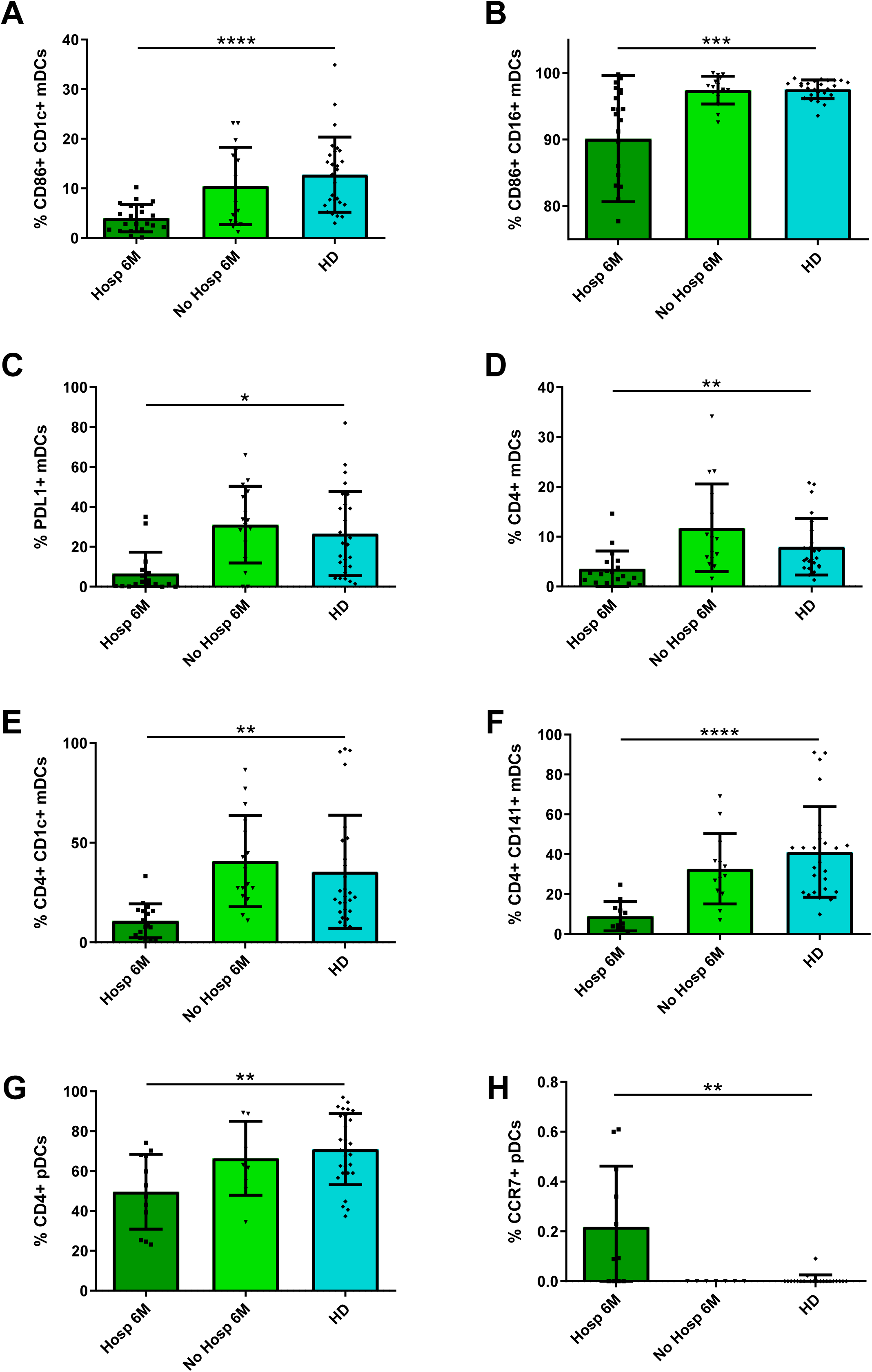
Some DC activation markers are not normalized in previously hospitalized patients seven months after SARS-CoV-2 infection. Bar graphs representing the percentage of DC subpopulations expressing CD86 (A - B), PD-L1 (C), CD4 (D - G) and CCR7 (H) in previously hospitalized (Hosp 6M) or previously non-hospitalized (No Hosp 6M) patients 6 months after SARS- CoV-2 infection and in healthy donors (HD). The median with the interquartile range is shown and each dot represents an individual. *p < 0.05, **p < 0.01, ***p < 0.001, ****p < 0.0001. Mann-Whitney U test was used for groups’ comparisons.

### Some DC activation markers are not normalized neither in previously hospitalized nor in non-hospitalized patients seven months after SARS-CoV-2 infection

Importantly, when we focused on the expression of other DC activation markers, we observed a lower percentage of β7+ cells in all mDCs and pDCs from both hospitalized and non-hospitalized patients after seven months of infection compared to HD (Figure 6A-E). The levels were also lower for IDO+ in total mDCs, CD1c+ and CD141+ mDCs and pDCs (Figure 6F-I). Lastly, we also found that both hospitalized and non-hospitalized patients seven months after infection showed lower percentages of CCR7+ and CD4+ cells within CD16+ mDCs and PD-L1+ cells within pDCs compared to HD (Figure 6J-L). In conclusion, we demonstrated that the alterations in integrin β7 and IDO, associated with migration and tolerance, are not restored to normal levels neither in previously hospitalized nor in non-hospitalized patients seven months after SARS-CoV-2 infection.

**Figure 6.**
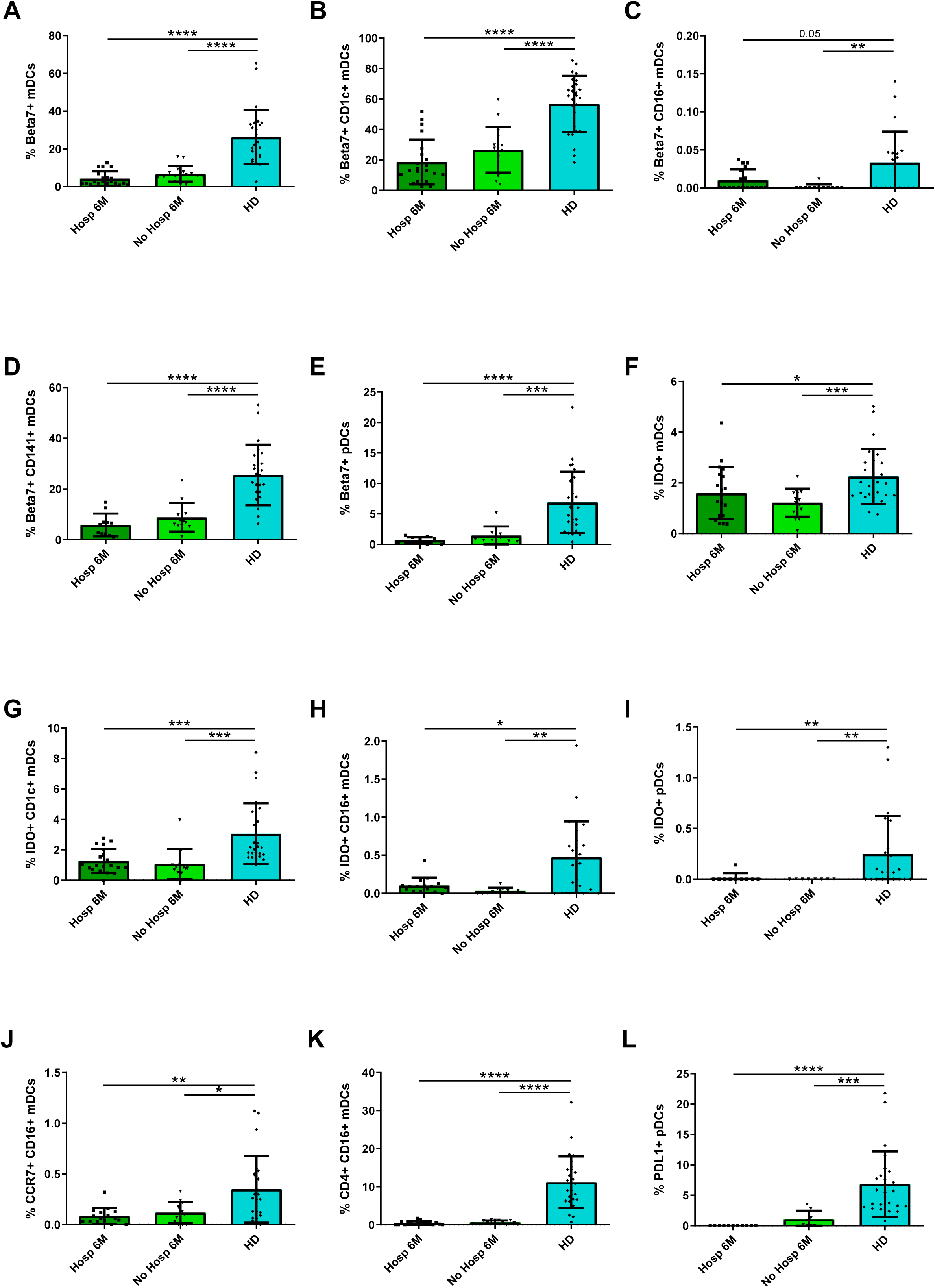
Some DC activation markers are not normalized neither in previously hospitalized nor in non-hospitalized patients seven months after SARS-CoV-2 infection. Bar graphs representing the percentage of DC subpopulations expressing β7 (A - E), IDO (F - I), CCR7 (J), CD4 (K) and PD-L1 (L) in previously hospitalized (Hosp 6M) or previously non-hospitalized (No Hosp 6M) patients 6 months after SARS- CoV-2 infection and in healthy donors (HD). The median with the interquartile range is shown and each dot represents an individual. *p < 0.05, **p < 0.01, ***p < 0.001, ****p < 0.0001. Mann-Whitney U test was used for groups’ comparisons.

## DISCUSION

The present study revealed that the deficits observed in CD1c+ mDCs and pDCs levels associated with altered homing and activation patterns in SARS-CoV-2 infected subjects in acute phase, were not restored beyond seven months after infection. Importantly, this long-term defects related to DC migration and tolerogenesis (integrin β7 and IDO expression) were present independently of whether or not the patients were previously hospitalized. In addition, hospitalized patients showed additional deficiencies related with DC activation.

pDCs are known to have an important role in the first line of defense against viral replication, which mainly resides in their capacity to produce IFN-I via TLR- 7/8 stimulation (Reizis, 2019). In this study, we first observed that acute SARS-CoV-2 infected patients displayed a dramatic decrease in pDC levels and a considerable reduction of IFN-α production. The strong direct correlation between pDC levels and IFN-α production suggested that this cell type was the main producer of this cytokine as it happens in other viral infections (Machmach et al., 2012). This reduction is in accordance with previous studies in animal models of SARS-CoV-1 infection, which associated this deficit with lethal pneumonia (Channappanavar et al., 2016) and is also consistent with recently published data following transcriptomic approaches (Hadjadj et al., 2020) and intracellular cytokine staining after TLR stimulation in SARS-CoV-2 infection (Arunachalam et al., 2020). Importantly, the low IFN-α production was the main parameter associated with disease severity, in agreement with previous studies (Arunachalam et al., 2020; Hadjadj et al., 2020), highlighting the potential use of this measurement as an early biomarker of disease progression. The mechanisms behind the attenuated IFN response have been related with viral antagonism of STAT1 (Signal transducer and activator of transcription 1) phosphorylation (Yang et al., 2020) and significantly, life-threatening ARDS in COVID-19 patients have been associated with neutralizing auto-antibodies against IFN-I (Bastard et al., 2020; Combes et al., 2021) and other inborn errors of IFN-I immunity (Zhang et al., 2020). These results support the essential role of IFN-I production in the first line of defense in COVID-19 for avoiding disease progression and point out to early immunotherapeutic strategies targeting this pathway.

Apart for IFN-I deficiency, one of the hallmarks of acute COVID-19 is the detection in plasma of heightened levels of soluble pro-inflammatory cytokines inducing a cytokine storm (Chen et al., 2020). Here, we found multiple correlations between DC numbers and activation markers with inflammatory marker and cytokine levels in acute SARS-CoV-2 infected patients. It was remarkable that the lower percentage of DCs was associated to higher levels of IL-6 and higher neutrophil numbers. High levels of IL-6 in COVID-19 patients have been widely related to a poorer disease progression (Laing et al., 2020). Moreover, neutrophils have been described as crucial drivers of hyperinflammation in COVID-19 (Parackova et al., 2020). It has to be also underlined, that the percentage of DCs expressing integrin β7 was inversely correlated to numerous inflammatory marker levels. These results suggest the hypothesis that not DC per se but DC migration to inflammatory sites may importantly contribute to the cytokine storm observed in SARS-CoV-2 infected patients. Our results also showed that patients with distinct level of disease severity displayed different associations of DC numbers and activation markers with inflammation. Therefore, DCs might be important contributors to the high inflammatory status characteristic of COVID-19 patients and this may dictate subsequent clinical progression.

The decreased numbers of total mDCs, CD1c+ mDCs and pDCs found in acute SARS-CoV-2 infected patients were in accordance with previous publications (Sánchez-Cerrillo et al., 2020; Zhou et al., 2020a). This fact might be explained by different mechanisms, including apoptosis due to increased inflammatory mediators produced by abortive SARS-CoV-2 infection of myeloid cells (Zheng et al., 2020). Another non-exclusive explanation could be that DCs migrate from peripheral blood to tissues or inflammatory sites, such as CD1c+ mDCs preferential migration to the lungs in patients with severe COVID-19 (Sánchez-Cerrillo et al., 2020). These defects were accompanied by alterations mainly found in activation, migration and tolerogenic markers that importantly persisted seven months after infection in previously hospitalized and also in non-hospitalized patients. Especially persistent in the total and DC subsets was the decreased expression of integrin β7. The expression of αEβ7 defines migration to antigen presentation sites within lymph nodes (Pribila et al., 2004) and α4β7 on mDCs and pDCs is indicative of migration of these cells to gut (Clahsen et al., 2015). Remarkably, SARS-CoV-2 has been shown to infect and productively replicate in human small intestinal organoids, increasing cytokine production and human angiotensin-converting enzyme 2 expression (Lamers et al., 2020). It has been also reported, that the disruption in gut barrier integrity contributes to COVID-19 severity (Giron et al., CROI 2021). Thus, the lower percentage of DCs expressing integrin β7 in peripheral blood might be a consequence of ongoing DC migration to the gut or other tissues or inflammatory sites up to seven months after infection. In fact, necropsy studies in SARS-CoV-2 infected patients have shown mononuclear inflammatory infiltrates in different organs (Xu et al., 2020). Also prominent was the deficit in IDO expressing DCs seven months after infection. In contrast, IDO+ CD1c+ and CD16+mDC levels in acute infection were dramatically increased compared to HD. This is in agreement with other acute respiratory infections such influenza (Fox et al., 2014) and respiratory syncytial virus (Ajamian et al., 2015) in which IDO expression is increased in order to counteract excessive inflammation as happen after acute SARS-CoV-2 infection. However, in this infection, the tissue damage in low respiratory tract is prominent (Broggi et al., 2020) and may persist at the long-term what may cause the exhaustion of IDO producing DCs and/or migration of these cells to inflammatory focus even after seven months after infection. Although these defects were present independently of whether or not the participants were previously at the hospital, hospitalized patients showed additional defects. These were, lower expression of the co-stimulatory molecule CD86, found in acute infection also by other authors (Arunachalam et al., 2020; Parackova et al., 2020; Zhou et al., 2020a) that persisted seven months after infection together with lower levels of CD4+ DCs. Low levels of activation molecules, such as CD86 have been related with a possible impairment in T cell and DCs response to the virus. Specifically, we and others have found pDC hypo-responsiveness to HIV after CD4 downregulation in this cell type (Barblu et al., 2012; Machmach et al., 2012). On the contrary, CCR7+ pDCs remained at high levels even after seven months after infection indicating again ongoing migration to lymph node or other inflammatory foci. In this line, the higher expression of other chemokine receptors such as CCR1, CCR3 and CCR5 has previously described in SARS-CoV-1 infected monocyte derived DCs (Law et al., 2009).

It is unknown whether these defects in the DCs compartment will be reversible after longer follow up or specific therapies may be needed for the normalization of these defects. What is clear is that persisting symptoms and unexpected substantial organ dysfunction are observed in an increasing number of patients who have recovered from COVID-19 (Carfì et al., 2020). Actually, Huang C et al. recently described that seven months after illness onset, 76% of the SARS-CoV-2 infected patients reported at least one symptom that persisted, being fatigue or muscle weakness the most frequently reported symptoms (Huang et al., 2021). In addition, many of those previously hospitalized patients presented residual chest imaging abnormalities, impaired pulmonary diffusion capacity and other extrapulmonary manifestations as a low estimated glomerular filtration rate (Huang et al., 2021). The immune mechanisms that might be involved in the development of these persisting symptoms are still unknown. However, it would be expected that seven months after SARS-CoV-2 infection there is still an inflammatory response due to persistent tissue damage or persistence presence of viral antigens in the absence of viral replication which may cause these deficits in DC. In fact, it has been reported that SARS-CoV-2 can persist in the intestines up to seven months following symptoms resolution (Tokuyama et al., CROI 2021). Thus, we postulate that the decrease in peripheral DCs numbers, along with the alterations in DC homing and activation markers seven months after the infection might be indicative of DC migration to inflammatory sites which may be contributing to long-term symptoms, a phenomenon also known as long COVID.

In summary, we have demonstrated that SARS-CoV-2 infected patients showed a deficit in some DC subsets and alterations in DC homing and activation markers, which are not restored more than seven months after the infection independently of previous hospitalization. Our results suggest that there is an ongoing inflammation which could be partially induced by DCs, these findings might contribute to a better understanding of the immunological sequelae of COVID-19.

## LIMITATIONS OF THE STUDY

All patients included in this study belong to the first wave of COVID-19 in Spain. It would have been interesting to have access to tissue samples, however due to safety issues at that moment of the pandemic it was not possible. At that time, different experimental treatments with very limited but transitory immunosuppressive effects were administered what may have affected the levels of immune parameters. However, the agreement of our observations with other data in the literature during acute infection and the persistence of these defects seven months after infection minimized the potential bias of these treatments in our results.

## MATERIAL AND METHODS

### EXPERIMENTAL MODELS AND SUBJECT DETAILS

#### HUMAN SUBJECTS / STUDY PARTICIPANTS

Seventy one participants with confirmed detection of SARS-CoV-2 by reverse-transcription polymerase chain reaction (RT-PCR) as previously described (To et al., 2020) were included. Out of these 71, 33 were hospitalized in acute phase of COVID-19 from March 25^th^ to May 8^th^ 2020, while 38 participants were recruited seven months after being diagnosed with COVID-19, from September 9^th^ to November 26^th^ 2020. These participants came from the COVID-19 patients’ Cohort Virgen del Rocio University Hospital, Seville (Spain) and the COVID-19 Cohort IIS Galicia Sur (CohVID GS), Vigo (Spain). Twenty-seven healthy donors (HD), with cryopreserved pre-COVID-19 samples (May 12^th^ to July 18^th^ 2014) were included from the HD cohort, collection of samples of the Laboratory of HIV infection, Andalusian Health Public System Biobank, Seville (Spain) (C330024). Written or oral informed consent was obtained from all participants. The study was approved by the Ethics Committee of the Virgen del Rocio University Hospital (protocol code “pDCOVID”; internal code 0896-N-20). Hospitalized participants during the acute phase of infection were divided in Mild (n=17) or Severe (n=16), based on the highest grade of disease severity during course of hospitalization. Severe participants were those who required Intensive Care Unit admission, or having ≥6 points in the score on ordinal scale based on Beigel et al. (Beigel New Engl J Med 2020) or death. Blood samples were collected at a median of 3 [interquartile range (IQR) 2 - 23] days after hospitalization and 14 [9 - 31] days after symptoms onset (Table S1). The group of participants discharged after infection, included previously hospitalized (n=21) and previously non hospitalized subjects (n=17). The samples from these participants were collected after a median of 201 [181 - 221] days after hospitalization and 208 [189 - 230] days after symptoms onset (Table S1). COVID-19 participants in the different groups were age and sex matched with HDs’ group (Table S1).

### STUDY DESIGN

Cross-sectional group comparison study. In a subgroup of participants, a longitudinal design was performed to study the kinetic of the studied parameters.

### METHODS DETAILS

#### PERIPHERAL BLOOD MONONUCLEAR CELLS (PBMCs) AND PLASMA ISOLATION

PBMCs from healthy donors and participants were isolated from fresh blood samples using BD Vacutainer® CPT™ Mononuclear Cell Preparation Tubes (with Sodium Heparin, BD Cat# 362780) in a density gradient centrifugation at the same day of blood collection. Afterwards, PBMCs were cryopreserved in freezing medium (90% of fetal bovine serum (FBS) + 10% dimethyl sulfoxide) in liquid nitrogen until further use. Plasma samples were obtained using BD Vacutainer™ PET EDTA Tubes centrifugation, aliquoted and cryopreserved at - 80°C until further use.

#### DENDRITIC CELLS IMMUNE PHENOTYPING

For dendritic cells (DCs) flow cytometry, PBMCs were centrifuged, pelleted and washed with Phosphate-buffered saline (PBS) and stained for 35 min at room temperature with LIVE/DEAD Fixable Aqua Dead Cell Stain (Life Technologies), BV421 CD86, BV650 CD11c, BV711 HLA-DR, BV786 CCR7 (CD197), FITC Lin-2 (CD3, CD14, CD19, CD20, CD56), BV605 CD16, PeCF594 PD-L1 (CD274), APC Integrin-β7 (BD Biosciences), PerCPCy5,5 CD4, APCCy7 CD1c, PeCy7 CD141 (BioLegend) and AF700 CD123 (R&D, San Diego, CA) antibodies. Then PBMCs were washed with Permeabilization Buffer 10X diluted 1:10 (eBioscience™), permeabilized by Fixation/Perm buffer (eBioscience™), and intracellularly stained with PE IDO (eBioscience, San Diego, CA, USA) antibody. DCs were gated based on Lin-2 HLA-DR expression. Each subset (mDCs and pDCS) was gated based on CD123 and CD11c expression. mDCs subsets were gated by using CD16, CD1c and CD141 staining, for gating strategy see Figure S1. Flow cytometry analyses were performed on an LRS Fortessa flow cytometer using FACS Diva software (BD Biosciences). Data were analyzed using the FlowJo software (Treestar, Ashland, OR). At least 1×10^6^ events were acquired per sample.

#### PBMCs CULTURE AND STIMULATION AND QUANTIFICATION OF IFN-α PRODUCTION

1×10^6^ thawed PBMCs were incubated at 37 ⁰C and 5% CO_2_ during 18 hours in RPMI with 10% FBS without any stimuli or with 1 µM CpG-A (ODN 2216; InvivoGen). After incubation, cells were pelleted and the supernatants conserved for the subsequent quantification of IFN-α production at -80°C. The amount of IFN-α in cell culture supernatants was assessed by an IFN-α multisubtype enzyme-linked immunosorbent assay kit (PBL Interferon Source Cat# 41105) according to the manufacturer’s instructions.

#### QUANTIFICATION OF CYTOKINE PRODUCTION IN PLASMA

Plasmas previously collected were used for the quantitative determination of cytokines. We used 3 different kits to quantify sCD25 by Human CD25/IL-2R alpha Quantikine ELISA Kit (R&D System, Cat# DR2A00), IP-10 by Human IP- 10 ELISA Kit (CXCL10) (Abcam, Cat# ab173194) and IL-6, IL-8, IL-1β, TNF-α, IFN-γ, MIP-1α, MIP-1β by MILLIPLEX MAP Human High Sensitivity T Cell Panel (Merck Cat# HSTCMAG-28SK) according to the manufacturer’s instructions.

### QUANTIFICATION AND STATISTICAL ANALYSIS

#### STATISTICAL ANALYSIS

Statistical analyses were performed by using Statistical Package for the Social Sciences software (SPSS 25.0; SPSS, Inc., Chicago, IL) and R environment 4.0.3 (2020-10-10), using RStudio Version 1.3.959 as the work interface and GraphPad Prism, version 8.0 (GraphPad Software, Inc.). ROUT method was utilized to identify and discard outliers. Differences between conditions among different groups were analyzed by two-tailed Mann-Whitney U test. The Wilcoxon test was used to analyze paired samples. The Spearman test was used to analyze correlations between variables. All differences with a P value of < 0.05 were considered statistically significant.

## ACKNOWLEDGMENTS

This study would not have been possible without the collaboration of Virgen del Rocio University Hospital COVID Team and COHVID-GS and IISGS Biobank. All of the patients, nursing staff, and data managers who have taken part in this project. We would like to thank to Sara Bachiller for the critical reading of the manuscript.

This work was supported by Consejeria de Transformacion Economica, Industria, Conocimiento y Universidades Junta de Andalucia (research Project CV20-85418), Consejeria de salud Junta de Andalucia (Research Contract RH- 0037-2020 to J.V.) the Instituto de Salud Carlos III (CP19/00159 to A.G.-V., FI17/00186 to M.R.J.-L., FI19/00083 to C.G-C, CM20/00243 to A.P-G and COV20/00698 to support COHVID-GS) and the Red Temática de Investigación Cooperativa en SIDA (RD16/0025/0020 and RD16/0025/0026), which is included in the Acción Estratégica en Salud, Plan Nacional de Investigación Científica, Desarrollo e Innovación Tecnológica, 2008 to 2011 and 2013 to 2016, Instituto de Salud Carlos III, Fondos FEDER. E.R.-M. was supported by the Spanish Research Council (CSIC).

Members of COHVID-GS (Galicia Sur Health Research Institute): Alexandre Araujo, Jorge Julio Cabrera, Víctor del Campo, Manuel Crespo, Alberto Fernández, Beatriz Gil de Araujo, Carlos Gómez, Virginia Leiro, María Rebeca Longueira, Ana López-Domínguez, José Ramón Lorenzo, María Marcos, Alexandre Pérez, María Teresa Pérez, Lucia Patiño, Sonia Pérez, Silvia Pérez-Fernández, Eva Poveda, Cristina Ramos, Benito Regueiro, Cristina Retresas, Tania Rivera, Olga Souto, Isabel Taboada, Susana Teijeira, María Torres, Vanesa Val, Irene Viéitez.

Members of the Virgen del Rocio University Hospital COVID Team: Jose Miguel Cisneros Herreros, Nuria Espinosa Aguilera, Cristina Roca Oporto, Julia Praena Segovia, José Molina, María Paniagua-García, Manuela Aguilar-Guisado, Almudena Aguilera, Clara Aguilera, Teresa Aldabo-Pallas, Verónica Alfaro-Lara, Cristina Amodeo, Javier Ampuero, Maribel Asensio, Bosco Barón-Franco, Lydia Barrera-Pulido, Rafael Bellido-Alba, Máximo Bernabeu-Wittel, Claudio Bueno, Candela Caballero-Eraso, Macarena Cabrera, Enrique Calderón, Jesús Carbajal-Guerrero, Manuela Cid-Cumplido, Juan Carlos Crespo, Yael Corcia- Palomo, Elisa Cordero, Juan Delgado, Alejandro Deniz, Reginal Dusseck- Brutus, Ana Escoresca Ortega, Fatima Espinosa, Michelle Espinoza, Carmen Ferrándiz-Millón, Marta Ferrer, Teresa Ferrer, Ignacio Gallego-Texeira, Rosa Gámez-Mancera, Emilio García, María Luisa Gascón-Castillo, Aurora González-Estrada, Demetrio González, Rocío GonzálezLeón, Carmen Grande- Cabrerizo, Sonia Gutiérrez, Carlos Hernández-Quiles, Concepción Herrera- Melero, Marta Herrero-Romero, Carmen Infante, Luis Jara, Carlos Jiménez- Juan, Silvia Jiménez-Jorge, Mercedes Jiménez-Sánchez, Julia LanserosTenllado, José María Lomas, Álvaro López, Carmina López, Isabel López, Luis F López-Cortés, Rafael Luque-Márquez, Daniel Macías-García, Luis Martín-Villén, Aurora Morillo, Dolores Nieto-Martín, Francisco Ortega, Amelia Peña-Rodríguez, Esther Pérez, Rafaela Ríos, Jesús F Rodríguez, María Jesús Rodríguez-Hernández, Santiago Rodríguez-Suárez, Ángel Rodríguez-Villodres, Nieves Romero-Rodríguez, Ricardo Ruiz, Zaida Ruiz de Azua, Celia Salamanca, Sonia Sánchez, Javier Sánchez-Céspedez, Victor Manuel Sández- Montagut, Alejandro Suárez Benjumea, and Javier Toral.

## AUTHOR CONTRIBUTIONS

P-G. A, V.J and G-C. C performed the experiments, analyzed and interpreted the data and participated in writing of the manuscript. G-V A, T-R M, S A, M-M E participated in data collection, data analysis and interpretation and performed experiments, J-L MR, R-I. B. M participated in manuscript data interpretation, R-J I, I. C, C. JC participated in data collection, S. C, R-O C, E. N, F-V. A, C. M. participated in data collection and manuscript interpretation. L.C. LF and E. V. participated in manuscript data analysis, patient and data collection, interpretation/ discussion of the results and coordination. E. R-M, participated in data analysis and interpretation, writing, conceived the idea and coordinate the project. P-G. A, V.J and G-C. C contributed equally to this work.

