## Supplemental information for "Dendritic cell deficiencies persist seven months after SARS-CoV-2 infection"

### Figure S1

**A**

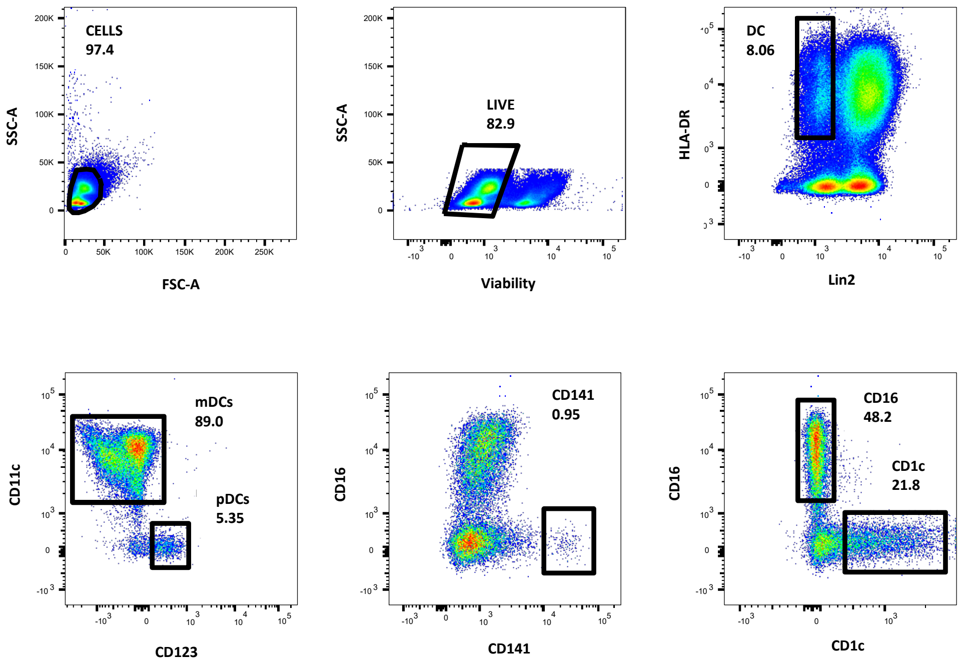
CELLS
97.4
SSC-A
FSC-A
LIVE
82.9
Viability
DC
8.06
HLA-DR
Lin2
mDCs
89.0
pDCs
5.35
CD11c
CD123
CD141
0.95
CD16
CD141
CD16
48.2
CD1c
21.8
CD1c

**B**

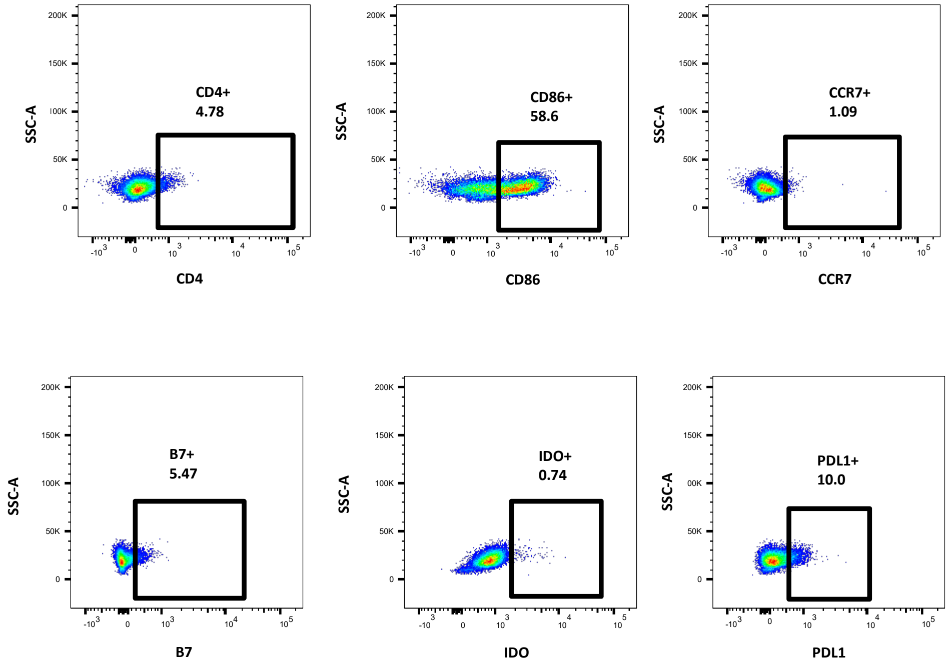
CD4+
4.78
SSC-A
CD4
CD86+
58.6
CD86
CCR7+
1.09
CCR7
B7+
5.47
B7
IDO+
0.74
IDO
PDL1+
10.0
PDL1

Figure S2

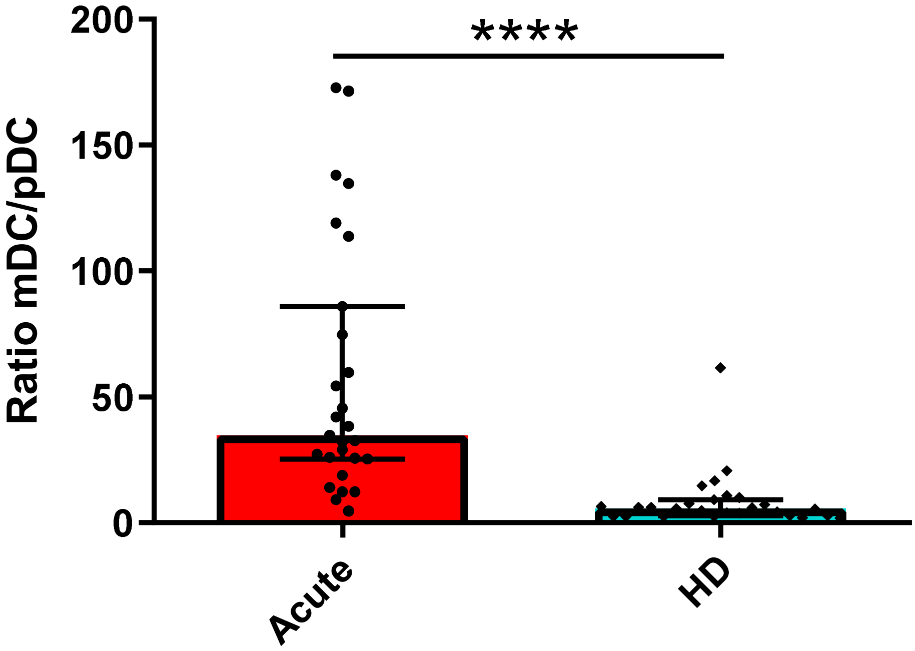

### Figure S3

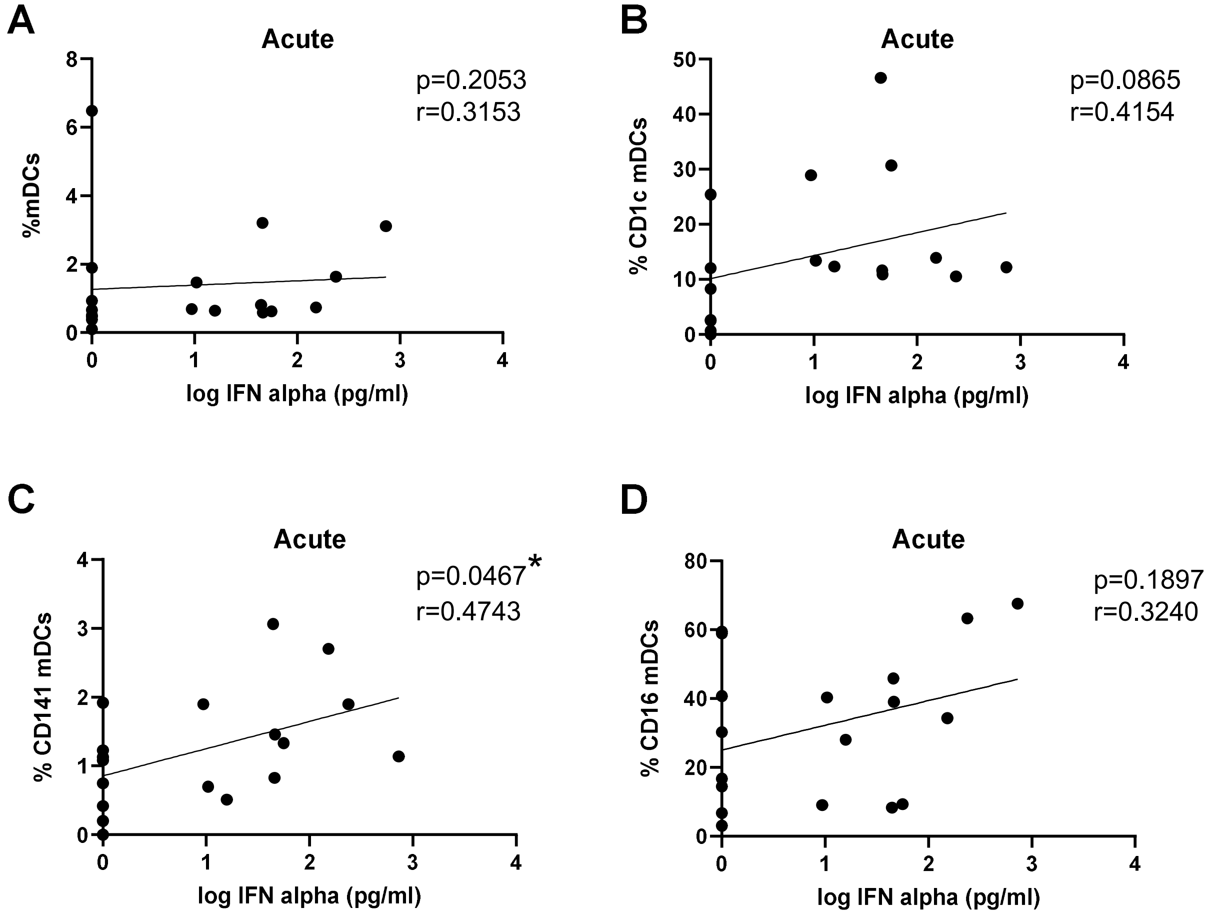
A
Acute
p=0.2053
r=0.3153
%mDCs
log IFN alpha (pg/ml)
B
Acute
p=0.0865
r=0.4154
% CD1c mDCs
log IFN alpha (pg/ml)
C
Acute
p=0.0467*
r=0.4743
% CD141 mDCs
log IFN alpha (pg/ml)
D
Acute
p=0.1897
r=0.3240
% CD16 mDCs
log IFN alpha (pg/ml)

Figure S4

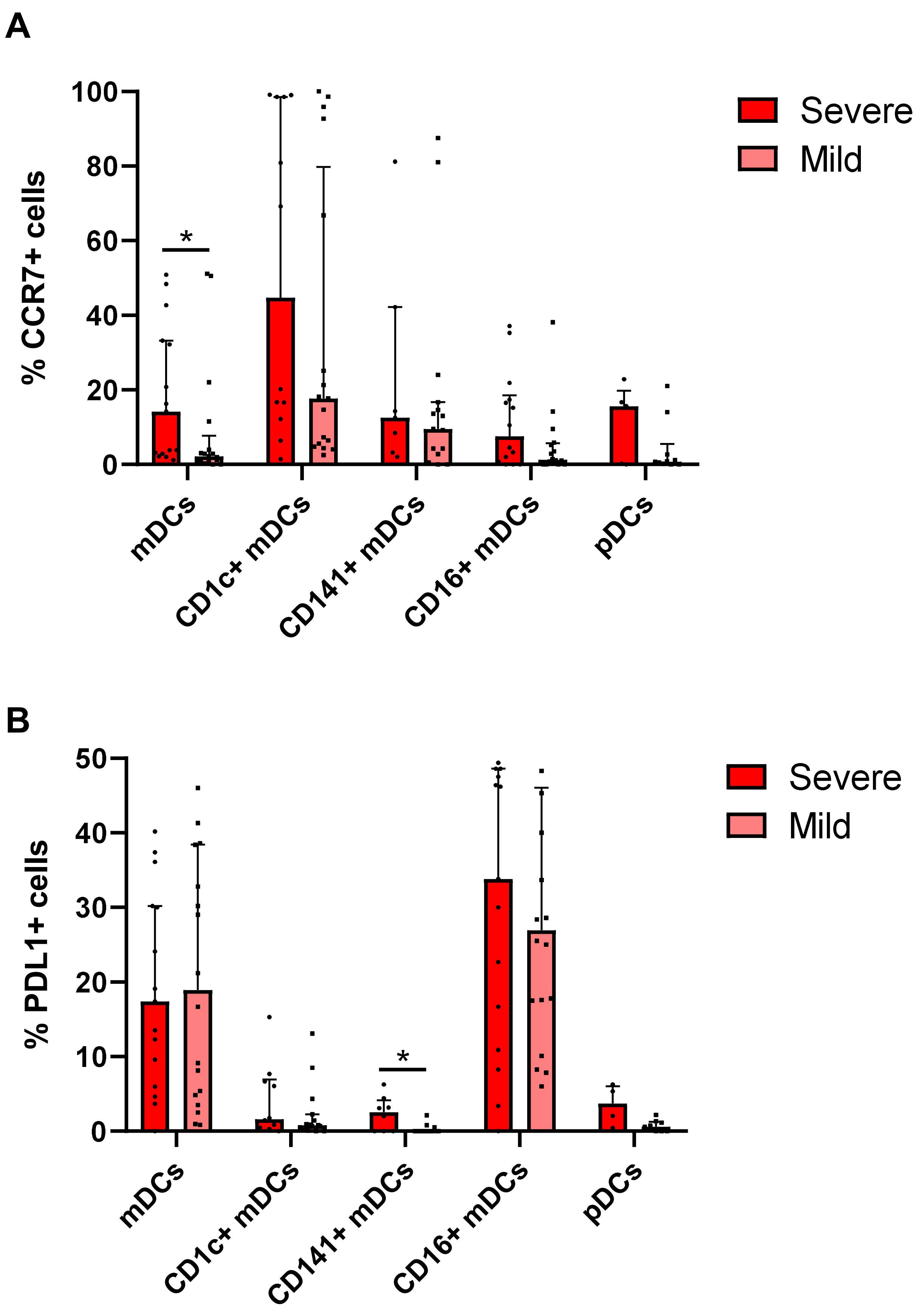
A
% CCR7+ cells
100
80
60
40
20
0
*
mDCs
CD1c+ mDCs
CD141+ mDCs
CD16+ mDCs
pDCs
Severe
Mild
B
% PDL1+ cells
50
40
30
20
10
0
*
mDCs
CD1c+ mDCs
CD141+ mDCs
CD16+ mDCs
pDCs
Severe
Mild

### Figure S5

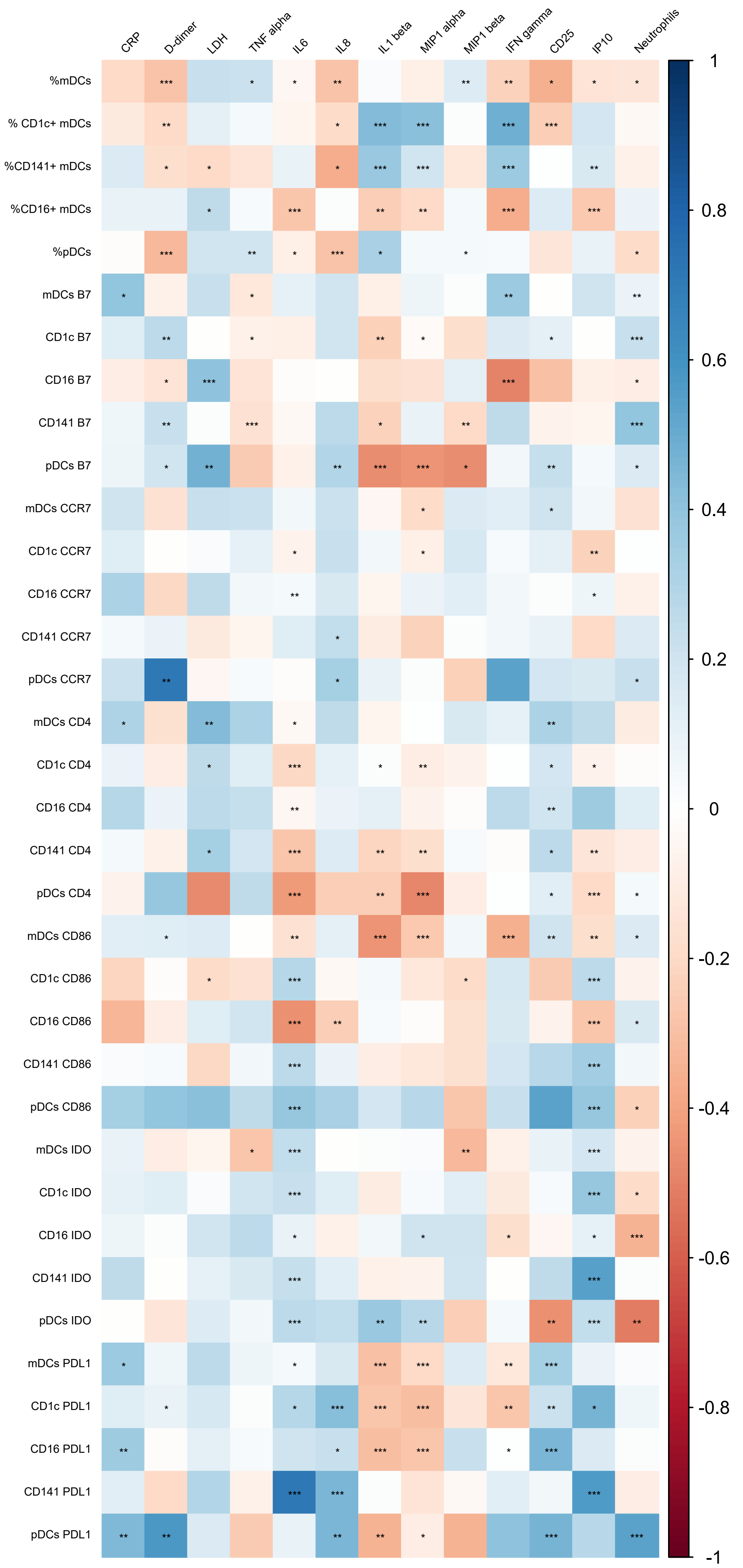
CRP
D-dimer
LDH
TNF alpha
IL6
IL8
IL1 beta
MIP1 alpha
MIP1 beta
IFN gamma
CD25
IP10
Neutrophils
%mDCs
% CD1c+ mDCs
%CD141+ mDCs
%CD16+ mDCs
%pDCs
mDCs B7
CD1c B7
CD16 B7
CD141 B7
pDCs B7
mDCs CCR7
CD1c CCR7
CD16 CCR7
CD141 CCR7
pDCs CCR7
mDCs CD4
CD1c CD4
CD16 CD4
CD141 CD4
pDCs CD4
mDCs CD86
CD1c CD86
CD16 CD86
CD141 CD86
pDCs CD86
mDCs IDO
CD1c IDO
CD16 IDO
CD141 IDO
pDCs IDO
mDCs PDL1
CD1c PDL1
CD16 PDL1
CD141 PDL1
pDCs PDL1
1
0.8
0.6
0.4
0.2
0
-0.2
-0.4
-0.6
-0.8
-1

Figure S6

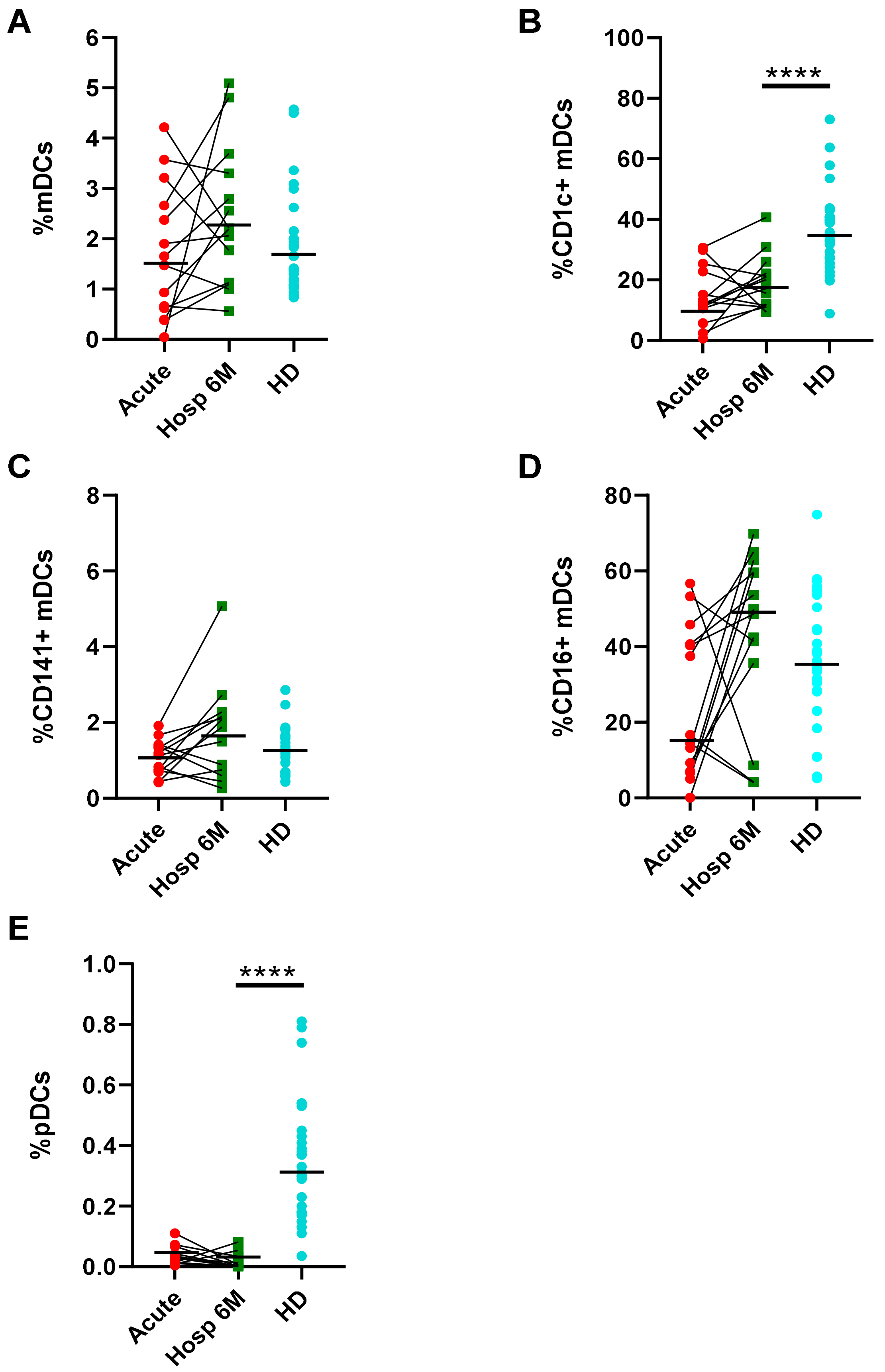

A
%mDCs
0
1
2
3
4
5
6
Acute
Hosp 6M
HD
B
%CD1c+ mDCs
0
20
40
60
80
100
****
Acute
Hosp 6M
HD
C
%CD141+ mDCs
0
2
4
6
8
Acute
Hosp 6M
HD
D
%CD16+ mDCs
0
20
40
60
80
Acute
Hosp 6M
HD
E
%pDCs
0.0
0.2
0.4
0.6
0.8
1.0
****
Acute
Hosp 6M
HD

### Figure S7

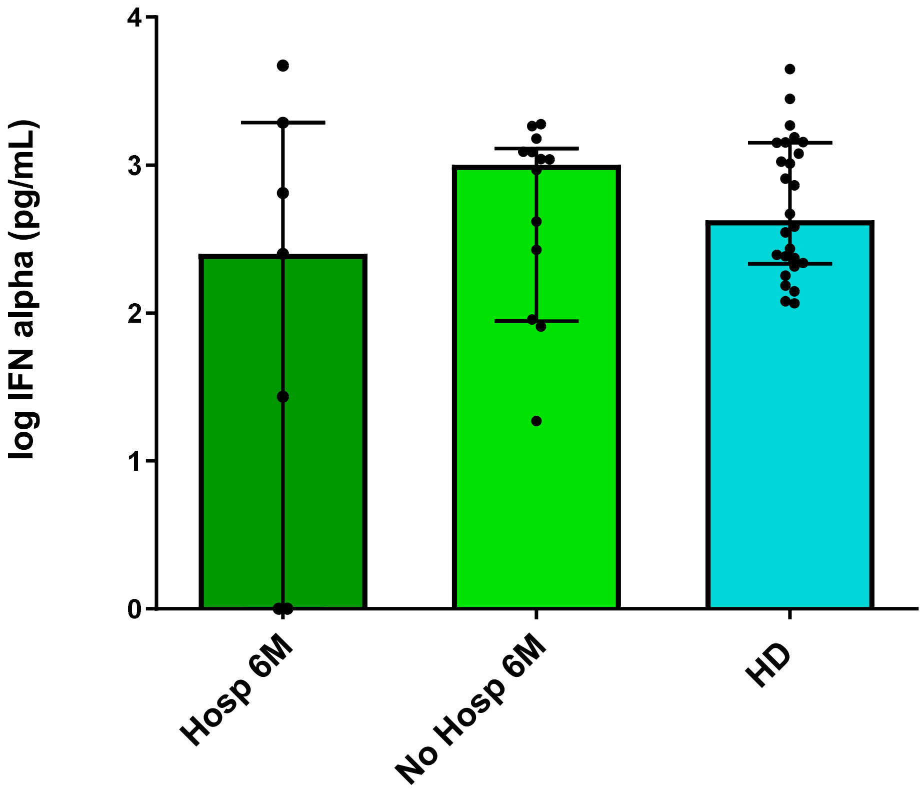

#### SUPPLEMENTAL INFORMATION

##### Figure S1. Gating strategy for the identification of DC subpopulations and activation markers

Pseudocolor dot plots showing the gating strategy used for the identification of studied populations in a representative SARS-CoV-2 infected patient. (A) Mononuclear cells were selected according to their size (FSC-A) and complexity (SSC-A) and dead cells were discarded using a viability marker. DC and subpopulations were identified as follows: total DCs (Lin2- HLA-DR+), total mDCs (CD11c+ CD123-), pDCs (CD11c- CD123+), and within mDCs: CD1c+, CD141+ and CD16+. (B) Representative data of total mDCs showing selected gates to analyze the percentages of CD4+, CD86+, CCR7+, β7+, IDO+ and PD-L1+ cells.

##### Figure S2. mDC/pDC ratio in acute SARS-CoV-2 infected patients and healthy donors

Bar graphs representing the ratio mDC/pDC in acute SARS-CoV-2 infected patients (acute) and healthy donors (HD). The median with the interquartile range is shown and each dot represents one individual. ****$p < 0.0001$. Mann-Whitney U test was used for groups' comparisons.

##### Figure S3. Association of mDC numbers with IFN-α production

Correlations between the percentages of total mDCs, CD1c+, CD141+ and CD16+ mDCs with IFN-α production in response to CpG-A in acute SARS-CoV-

2 infected patients (A-D). Each dot represents an individual. *p < 0.05. Spearman test was used for non-parametric correlations.

**Figure S4. DC markers expression in SARS-CoV-2 infected patients with severe and mild symptoms.**

Bar graphs representing the percentage of DCs expressing CCR7 (A) and PD-L1 (B) in acute severe and mild SARS-CoV-2 infected patients. The median with the interquartile range is shown and each dot represents an individual. *p < 0.05. Mann-Whitney U test was used for groups' comparisons.

**Figure S5. Associations of DC numbers and activation markers with inflammatory markers in acute SARS-CoV-2 infected patients**

Heatmap graphs representing correlations between the percentages of DC subpopulations and the percentages of DCs expressing activation and homing markers with inflammatory markers including CRP, D-dimer, LDH, TNF-α, IL-6, IL-8, IL1-β, MIP1-α, MIP1-β, IFN-γ, sCD25, IP-10 and neutrophil numbers, in acute SARS-CoV-2 infected patients. Blue color represents positive correlations and red color shows negative correlations. The intensity of the color indicates the R coefficient. *p < 0.05, **p < 0.01, ***p < 0.001. Spearman test was used for non-parametric correlations.

**Figure S6. Paired analysis of DC subsets of SARS-CoV-2 infected patients in acute phase and seven months after the infection**

Before and after graphs representing the paired analysis of the percentage of total mDCs, CD1c+, CD141+ and CD16 mDCs and pDCs (A - E) in patients in acute phase (Acute) and seven months after SARS-CoV-2 infection (Hosp 6M) and in healthy donors (HD). The median is shown and each dot represents an individual. ****$p < 0.0001$. Wilcoxon test was used for paired samples and Mann-Whitney U test was used for groups' comparisons.

**Figure S7. IFN-α production seven months after SARS-CoV-2 infection**

Bar graphs representing the IFN-α production of PBMCs in response to CpG-A in previously hospitalized (Hosp 6M) or previously non-hospitalized (No Hosp 6M) patients seven months after SARS-CoV-2 infection and in healthy donors (HD). The median with the interquartile range is shown and each dot represents an individual. Mann-Whitney U test was used for groups' comparisons.

Supplementary table 1. Characteristics of the study patients.

| | **Acute Infection** | | | **Discharged (6-8 months after diagnosis)** | | | **Healthy Donors** |
|---|---|---|---|---|---|---|---|
| | **All (n=33)** | **Mild (n=17)** | **Severe (n=16)** | **All (n=38)** | **Previously Hospitalized (n=21)** | **Previously Non Hospitalized (n=17)** | **(n=27)** |
| Age (years) | 66 [59-77] | 62 [57-78] | 69 [63 – 73] | 67 [60 – 72] | 68 [63 – 73] | 65 [58 – 71] | 62 [39 – 84] |
| Sex (Female sex), n (%) | 12 (36) | 7 (41) | 5 (31) | 17 (48) | 7 (33) | 10 (59) | 11 (41) |
| Oxygen Saturation ($SatO_2$), (%) | 95 [91 – 98] | 96 [95 – 99] | 92 [90 – 95] | N/A | N/A | N/A | N/A |
| Time since hospitalization, (days) | 3 [2 – 23] | 2 [1 – 3] | 20 [3 – 31] | 201 [181 – 221] | 183 [168 – 197] | 221 [219 – 228] | N/A |
| Time since symptoms onset, (days) | 14 [9 – 31] | 11 [5 – 14] | 31 [19 – 38] | 208 [189 – 230] | 192 [179 – 203] | 230 [224 – 235] | N/A |
| Time hospitalized, (days) | 16 [7 – 34] | 7 [5 – 10] | 28 [20 – 43] | N/A | 16 [8 – 40] | 0 | N/A |
| Comorbidities, n (%) | 26 (79) | 13 (77) | 13 (81) | 19 (50) | 16 (76) | 3 (18) | N/A |
| Diabetes mellitus | 8 (24) | 4 (24) | 4 (25) | 6 (16) | 5 (24) | 1 (6) | N/A |
| Hypertension | 19 (57) | 8 (47) | 11 (70) | 15 (40) | 13 (62) | 2 (12) | N/A |
| Cardiovascular disease | 7 (21) | 4 (24) | 3 (19) | 8 (21) | 6 (29) | 2 (12) | N/A |
| Obstructive pulmonary disease | 5 (15) | 3 (18) | 2 (13) | 4 (11) | 4 (19) | 0 | N/A |
| Malignancy | 2 (6) | 1 (6) | 1 (6) | 2 (5) | 2 (10) | 0 | N/A |
| Symptoms at admission (%) | | | | | | | |
| Cough | 20 (61) | 10 (59) | 10 (63) | 29 (76) | 19 (91) | 10 (59) | N/A |
| Fever | 22 (67) | 10 (59) | 12 (75) | 28 (74) | 16 (76) | 12 (71) | N/A |
| Dyspnea | 14 (43) | 7 (41) | 7 (44) | 21 (55) | 15 (71) | 6 (35) | N/A |
| Anosmia | 6 (18) | 3 (18) | 3 (19) | 4 (11) | 4 (19) | 0 | N/A |
| Diarrhoea | 7 (21) | 4 (24) | 3 (19) | 12 (32) | 9 (43) | 3 (18) | N/A |
| Muscle pain | 6 (18) | 2 (12) | 4 (25) | 4 (11) | 3 (14) | 1 (6) | N/A |
| Treatment during hospitalization; n (%) | | | | | | | |
| Hydroxychloroquine | 28 (85) | 13 (77) | 15 (94) | N/A | 20 (95) | N/A | N/A |
| Lopinavir/Ritonavir | 20 (61) | 7 (41) | 13 (81) | N/A | 15 (71) | N/A | N/A |
| Beta Interferon | 10 (30) | 1 (6) | 9 (56) | N/A | 9 (43) | N/A | N/A |
| Corticoids | 16 (49) | 4 (24) | 12 (75) | N/A | 8 (38) | N/A | N/A |
| Remdesivir | 3 (9) | 3 (18) | 0 | N/A | 0 | N/A | N/A |

| | | | | | | | |
|---|---|---|---|---|---|---|---|
| Tocilizumab | 9 (27) | 0 | 9 (56) | N/A | 7 (33) | N/A | N/A |

[a]Categorical variables are expressed as number and percentages (%), and continuous variables are expressed as median (interquartile ranges [IQR]). N/A, not applicable. The different groups (acute infection, discharged patients and healthy donors) were age and sex matched. Chi-square test and a Mann-Whitney U test were used to compare categorical and continuous variables, respectively. Analysis by age; acute infection vs HD (p=0.259); discharged patients vs HD (p=0.440); Previously Hospitalized patients vs HD (p=0.488); Previously Non Hospitalized patients vs HD (p=0.604). Analysis by sex; acute infection vs HD (p=0.793); discharged patients vs HD (p=0.803); Previously Hospitalized patients vs HD (p=0.765); Previously Non Hospitalized patients vs HD (p=0.354).
